# Targeting the untargeted: Uncovering the chemical complexity of root exudates

**DOI:** 10.1101/2024.09.17.613458

**Authors:** Katrin Möller, Annalena Ritter, Phillip J. Stobinsky, Kai Jensen, Ina C. Meier, Harihar Jaishree Subrahmaniam

## Abstract

The chemical complexity of root exudates has garnered significant attention in recent years, yet critical gaps remain in understanding the full scope of root exudate chemical variation across the plant kingdom. To address this, we conducted a systematic review of 57 studies, comprising 124 experiments, aimed at evaluating current methodologies and findings in untargeted root exudate chemical analysis. Our review revealed that hydroponic (44%) and soil-hydroponic hybrid (32%) sampling approaches, primarily utilising water as the collection medium, were the most common experimental setups. Liquid chromatography-mass spectrometry (LC-MS) was the predominant analytical technique used in 54% of the studies, followed by gas chromatography-mass spectrometry (GC-MS) in 31%. The average number of metabolites identified per analysis was 960, though the number of annotated metabolites varied considerably. Shikimates, phenylpropanoids, and carbohydrates were the most frequently identified classes, with their relative abundances varying widely. Several methodological challenges were highlighted, including inconsistencies in sampling techniques, underrepresentation of non-crop plants, and incomplete chemical annotation. To address these limitations, we propose a framework emphasising the need for representative exudate sampling, the use of multiple analytical approaches, the development of advanced bioinformatics tools, and the integration of these findings to enhance our understanding of root exudates and their ecological functions.

## Introduction

The rhizosphere is the main interface for plant-soil interactions and a hotspot of microbial activity and biogeochemical processes. Despite its narrow extension adjacent to roots, the rhizosphere has the potential to influence large-scale biogeochemical cycles (Guenet et al., 2010; Philippot et al., 2013). A major mechanism for plants to mediate microbial activities in the rhizosphere is *via* the release of metabolites by root exudation, collectively called the exudate metabolome. This metabolome comprises a diverse array of low molecular weight (MW: <1000 Da) and high molecular weight (MW: >1000 Da) organic and inorganic compounds (Oburger & Jones, 2018). Organic compounds exuded include primary and secondary metabolites from different chemical compound classes like (i) carbohydrates, organic acids, amino acids and fatty acids (primary metabolites) and (ii) alkaloids, phenolics and terpenoids (secondary metabolites) (Badri & Vivanco, 2009). The quantity and chemical composition of root exudates can vary strongly depending on plant species, plant age, root morphology (including root length, branching, and root hair density) (Gao et al., 2024; Staszel-Szlachta et al., 2024), interactions with plant neighbours and rhizosphere microbes, as well as climatic conditions and nutrient availability, among others (Badri & Vivanco, 2009; Sasse et al., 2018; van Dam & Bouwmeester, 2016).

A large variety of experimental designs have been developed to sample root exudates (see detailed reviews by Oburger and Jones (2018) or Salem et al. (2022)). Different sampling approaches address the growth conditions of plants with a variation of sampling durations and media to collect the exudates. Hydroponic approaches, including pure hydroponics, semi-hydroponics, and aeroponics, offer controlled environments with plants growing in different solutions to collect exudates directly, minimising soil contamination and microbial interference (Lin et al., 2023; Maver et al., 2022). Soil- hydroponic hybrid systems combine soil-grown plant roots with hydroponic setups to capture exudates under more natural conditions (Dietz et al., 2019; Herz et al., 2018). Sorption-based approaches utilise resins, filter papers, agar sheets or other sorptive materials to capture exudates *in situ*, offering a semi-quantitative analysis while accounting for potential interactions with the soil matrix and microbial activity (Asaduzzaman et al., 2014; Shi et al., 2023). Wu et al. (2000) also introduced a method of growing plants directly on agar from which metabolites can be extracted. Rarely, root exudates have been sampled exclusively soil-based, by extracting them directly from soil (Li et al., 2022). Sampling media to collect exudates are mostly some form of aqueous (e.g. distilled, double-distilled, ultrapure water) (Döll et al., 2024) or nutrient solutions (McLaughlin et al., 2023; Phillips et al., 2008), but also alcoholic solutions (e.g. methanol or ethanol) (Sorty et al., 2023) or ionic solutions (e.g. calcium chloride) (Fortier et al., 2023) are used. The sampling duration typically ranges from minutes to hours but can span multiple days to weeks (Alahmad et al., 2024; Asaduzzaman et al., 2014; Zubair et al., 2017). This is a crucial factor, as prolonged durations can induce hypoxic stress (Jackson & Armstrong, 1999) or result in reuptake of exudates (Farrar et al., 2003; Tiziani et al., 2020), while short durations may fail to capture trace compounds (Alahmad et al., 2024; Oburger & Jones, 2018).

The chemical annotation of root exudate metabolites is essential to understand their functional significance in shaping biogeochemical feedback-loops, influencing soil physical properties and mediating biological interactions within the rhizosphere. Because there is a wide variety of plant metabolites, they need to be grouped in a context-relevant way. Novel natural product (NP) classification algorithms offer a method of grouping metabolites by their specialised metabolism (pathway), chemical properties or chemotaxonomic information (superclass), and structural details (class) (Kim et al., 2021). Compound classes from this system, such as carbohydrates, fatty acids, amino acids, alkaloids, shikimates and phenylpropanoids, and terpenoids, provide a useful framework for describing the metabolomic functionality of root exudate metabolites.

Primary metabolites are generally released into the rhizosphere in larger quantities than secondary metabolites (Badri & Vivanco, 2009). Simple sugars such as glucose and fructose provide energy-rich substrates for soil microorganisms. Their release can stimulate microbial metabolisms, leading to the decomposition of organic matter and the mineralisation of key nutrients such as nitrogen and phosphorus (Bais et al., 2006; Philippot et al., 2009). Amino acids are also energy and nitrogen sources for soil fauna and can act as chemo-attractors for microorganisms (Allard-Massicotte et al., 2016). Fatty acids are an important energy source for mycorrhizal fungi in their symbiosis with plants (Jiang et al., 2017). While less abundant, secondary metabolites perform highly specialised and critical ecological functions within the rhizosphere, especially in plant defence mechanisms. Terpenoids, such as saponins and sesquiterpenes, show strong antimicrobial characteristics (Dixon, 2001). Certain flavonoids act as signalling molecules that promote symbiotic relationships with arbuscular mycorrhizal (AM) fungi or rhizobia, thereby enhancing nutrient uptake and nitrogen fixation (Zwetsloot et al., 2020). Alkaloids, such as nicotine and morphine, are another important group of secondary metabolites known for their potent bioactive properties, including anti-herbivory and insecticidal effects (Li et al., 2009). The shikimate pathway produces important secondary metabolites like tannins and lignins, which contribute to structural integrity and defence while also producing precursors for phenylpropanoids (Samal et al., 2023). Phenylpropanoids, such as cinnamic acid derivatives, have antimicrobial properties and are a main component of waxes in roots, forming a physical barrier against invading organisms (Döll et al., 2021).

One major challenge in understanding the complexity of root exudates remains prominent: the accurate analysis and chemical annotation of metabolites. Most studies have traditionally focused on targeted approaches (Oburger & Jones, 2018), where the analysis is directed towards specific, known metabolites within the exudates. These approaches involve meticulous sample preparation, often chromatography (IC), gas and liquid chromatography coupled with mass spectrometry (GC-MS and LC-MS), nuclear magnetic resonance spectroscopy (NMR) (Fortier et al., 2023) and fourier-transform ion cyclotron resonance mass spectrometry (FT-ICR-MS) (Calabrese et al., 2023; Miao et al., 2020; Shulaev, 2006). These targeted methods enable detailed investigation of specific metabolites under various environmental conditions, providing valuable insights into plant-species-specific exudation patterns (Carvalhais et al., 2011; Sandnes et al., 2005). However, because of this specific focus, they tend to overlook many metabolites that may play crucial roles in plant-soil interactions. This has left many exudate metabolites still uncharted. This gap in knowledge limits our ability to fully understand the vast biochemical diversity that plants release into the rhizosphere and the roles these compounds play in shaping plant-soil interactions. Continued advancements in metabolomic techniques, alongside a deeper exploration of the chemical and ecological complexity of exudates, are essential to bridging this knowledge gap. Here, untargeted approaches, like metabolomic fingerprinting, offer a broader perspective by aiming to capture the full diversity of metabolites released by roots (Macel et al., 2014; Weinhold et al., 2022). Recent advances in analytical instrumentation, particularly in mass spectrometry (MS), have facilitated these untargeted analyses, enabling the detection of a wide range of metabolites with high sensitivity (Salem et al., 2022). Despite their potential, untargeted approaches still face challenges, including the time-consuming nature of data analysis and metabolite annotation. However, new machine-learning approaches can help handle large volumes of data from metabolomic analyses in the future (Vlaar et al., 2022).

In this synthesis, we aimed to compile the status quo of methodological approaches in untargeted metabolomics and the metabolic composition of plant root exudates as analysed by different means. Our synthesis is meant to help inform researchers about the metabolic diversity and compound classes expected in root exudates. Future research may use this knowledge to target specific functions of single exudate compounds, combinations of different exuded compounds, or metabolic diversity of root exudation.

## Methods

A literature search was conducted to identify studies that analysed root exudates from different plant species and orders across various ecosystems by metabolomics using an untargeted approach. We used a range of keyword combinations to guide our search. Core search terms included ‘root exudates’, ‘root exudation’, ‘chemical composition’, and ‘untargeted’. We also explored variations such as ‘root exudate chemicals’, ‘untargeted profiling’, ‘metabolic profiling’, and ‘ecosystems’, applying Boolean operators to refine the results. To identify relevant studies, these combinations were systematically applied across selected databases, including Web of Science, Google Scholar, Semantic Scholar, and the Researcher app. Thereby, we gathered 57 studies in total, which were published between 2003 and 2024. No studies were found from earlier years that fulfilled our filtering criteria. Collected data from the studies include the plant species identity, root exudate sampling approach, medium and duration, method of metabolite analysis, total number of reported metabolites, and the total number of annotated metabolites (Supplementary Table 1). When a study performed different experiments, e.g. with multiple plant species, sampling durations or sampling media, information was given for each experiment, regardless of whether they were analysed separately or together. This resulted in 124 experiments in total. If a study performed multiple analyses (GC-MS, LC-MS, or other), we included them as separate analyses within the experiments, resulting in 109 analyses. The discrepancy between experiments and analyses results from studies where multiple experiments were conducted, but the metabolite analyses were reported together. We also calculated the percentage of annotated compounds from the total number of reported compounds. The metabolite data was obtained either from tables in the supplementary material of the selected papers, text, or figures within the papers. For studies in which treatments were applied (e.g. varying nutrient concentrations or infection with specific bacteria), only data from the control was included in our analysis. The category ‘crops’ contains all plants cultivated by humans and used for food, fodder, medicine, fibre or drugs. The category ‘model plants’ includes extensively studied plants which do not belong to the category of crops and are only studied because they are advantageous due to, e.g. a small genome or easy cultivation. The remaining trees, wetland plants, aquatic plants and forbs were classified as ’other plants’. The root exudate sampling approach was categorised as hydroponic, soil-hydroponic-hybrid, soil-extraction, agar, sorption, or aeroponic. For the sampling medium, we summarised ultrapure, sterile, deionised, distilled, and sterile distilled water into the category water. The method for metabolite analysis was either GC-MS, LC-MS, mixed (if more than one method was used), or other, which contains FT-ICR-MS, thin-layer chromatography (TLC), and NMR. As different studies used different ways of classification of chemical compounds, we decided to use the natural product classification (Dührkop et al., 2013; Kim et al., 2021), a relatively new and coarse classification system. We categorised the annotated compounds into the following categories: carbohydrates, fatty acids, amino acids and peptides, terpenoids, shikimates and phenylpropanoids, alkaloids, and polyketides. Every compound which did not fit into these categories was classified as other, such as organic acids (excluding fatty acids and amino acids), alcohols, vitamins, benzene derivatives, lignin, and nitriles.

## Results

### Exudate sampling techniques

There is a noticeable increase in interest in untargeted analysis of the root exudate metabolome (Figure 1). While only six studies on this topic were published between 2003 and 2014, fifteen studies were published between 2015 and 2019, and 36 studies from 2022 to May 2024. The increase in metabolomic studies on exudation is steeper than and delayed to the general increase of publications on plant ecology, with an exponential increase since about 2015. While this suggests that the interest in untargeted metabolomics is generally increasing, not all plant orders and species were studied equally.

**Figure 1:**
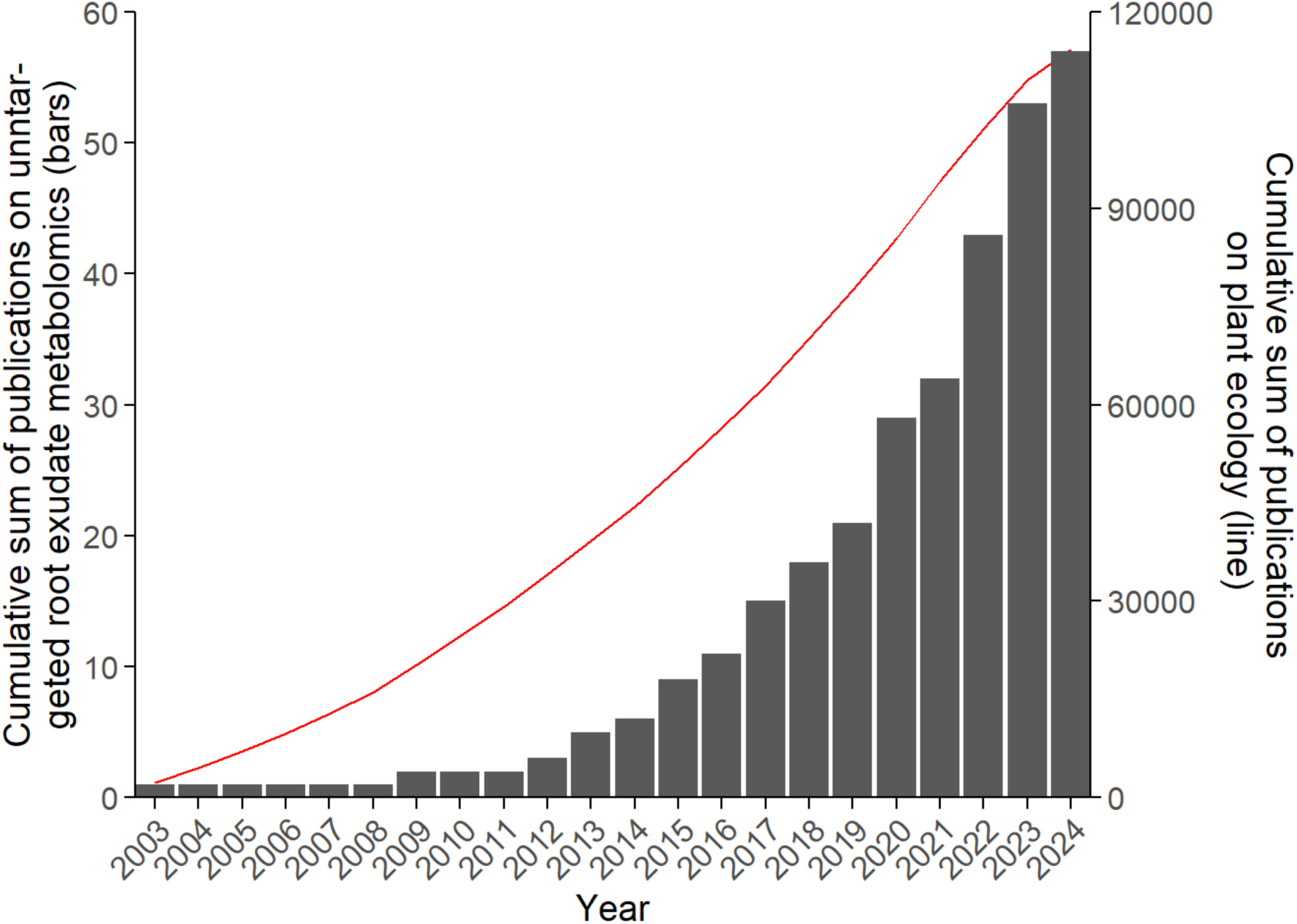
Cumulative sum of publications on untargeted root exudate metabolomics (bars) in relation to the cumulative sum of publications on plant ecology in general (red line). We gathered 57 studies in total, which were published between 2003 and 2024. No studies from earlier years fulfilled our filtering criteria.

The plats utilisation significantly influenced the number of experiments conducted (Figure 2). Nearly three-quarters of all experiments were conducted on crops, and only about a tenth on model plants. Within the crops, the plant order Poales accounts for almost half of all experiments, mainly due to many studies on *Zea mays* (a quarter of all crop studies). Fabales and Solanales together account for another quarter of the experiments on crops. The main model plant investigated for untargeted metabolomic analysis of root exudates is *Arabidopsis thaliana* (67% of experiments on model plants) from the plant order Brassicales. Only two model plants other than *Arabidopsis* were investigated (*Brachypodium distachyon* and *Medicago trunculata*). Among plants categorised differently from crops or model species, Coniferales represent approximately 30%.

**Figure 2:**
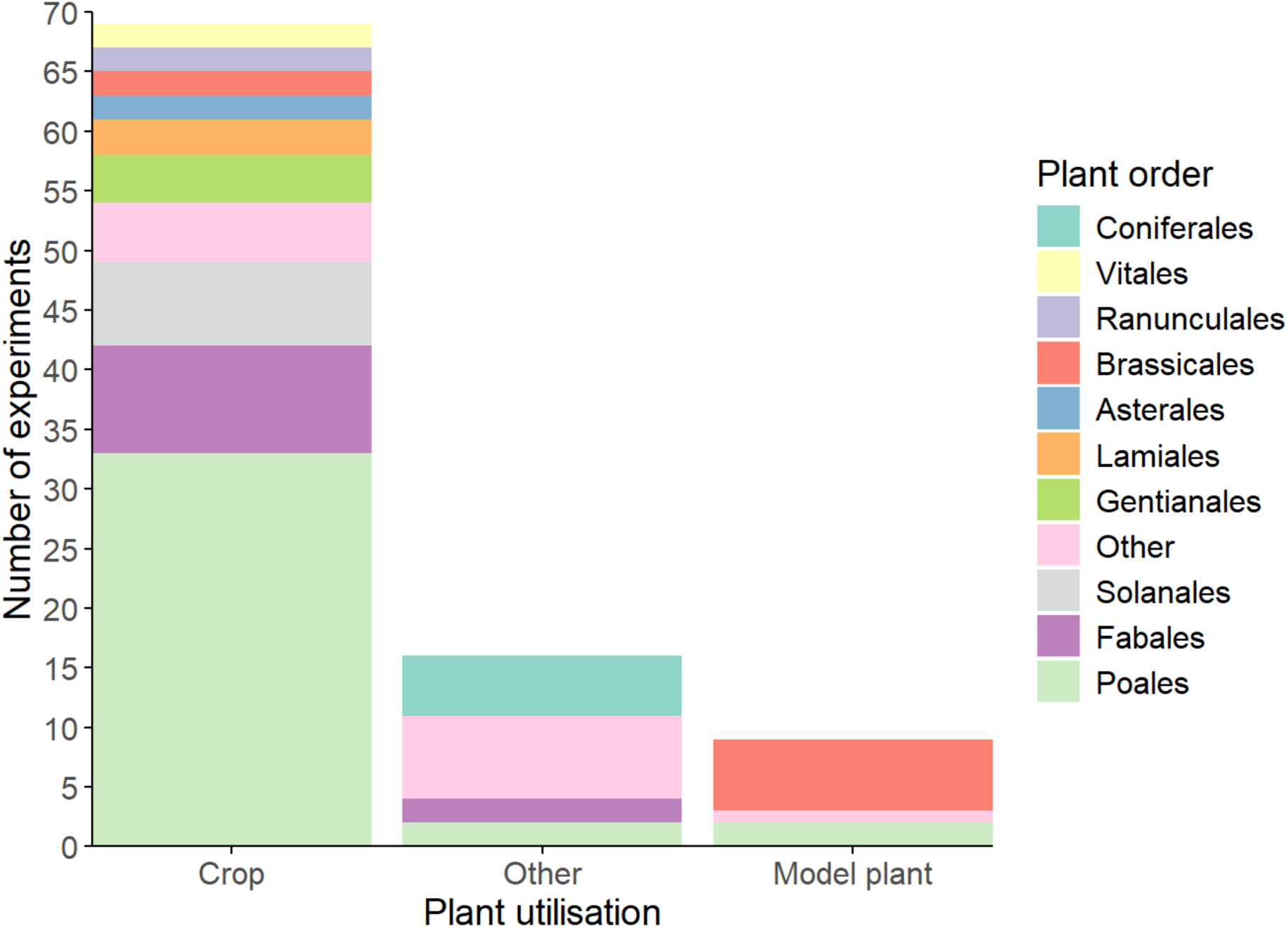
Contribution of different plant orders to the total number of experiments grouped by their targeted use. Plants not used as crops or model plants were grouped into ‘Other’. The plant order ‘Other’ includes all plant orders that only appear once in a plant utilisation group (Supplementary Table 2).

Various sampling approaches, media and durations were used in untargeted exudate metabolite experiments. Three-quarters of all studies used either hydroponic (44%) or soil-hydroponic-hybrid (32%) approaches utilising water (59 and 70%, respectively) or nutrient solutions (22 and 20%, respectively) for collecting root exudates (Figure 3). Only 15% of all studies used soil-extraction, often with methanol as the sampling medium (56%). Even fewer studies have used other methods, such as exudate collection in agar, via sorption, or with aeroponics, which comprise 10% of the total.

**Figure 3:**
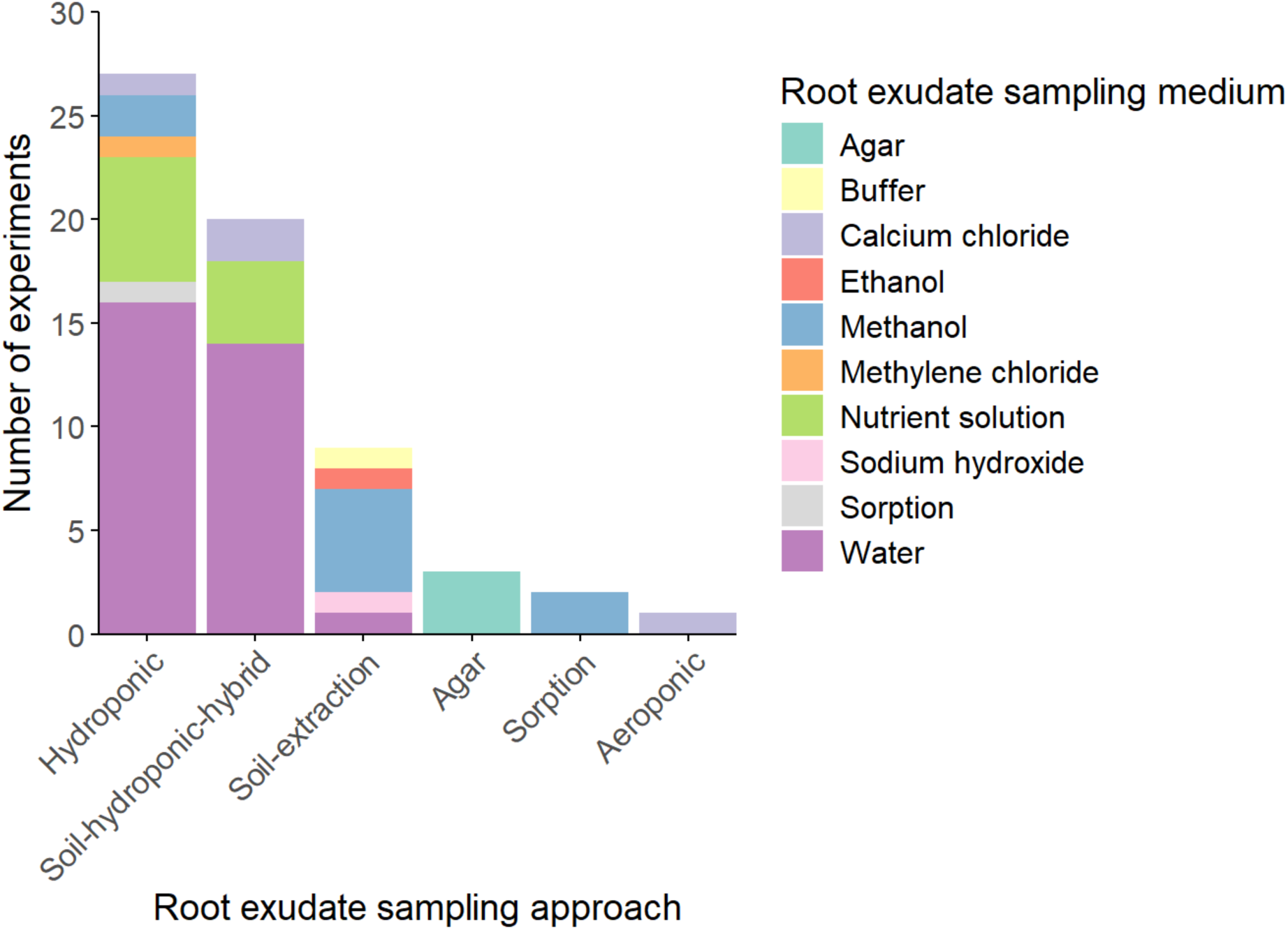
Contribution of different root exudate sampling approaches and sampling media to the total number of experiments. The ‘water’ category includes ultrapure, sterile, deionised, distilled, and sterile distilled water.

The sampling duration for trapping root exudates in the respective sampling medium spanned a wide range from less than one second to 56 days, with the most common duration being two hours, followed by two days and by six hours (Figure 4). Only 5% of the studies collected exudates in less than one hour. The maximum sampling duration for hydroponic-soil-hybrid was seven days, while the maximum for hydroponics was 45 days. Sampling for extraction from agar or soil always spanned longer than one day. The sampling duration is sometimes undefinable for soil-extraction because the plants grew in the soil for an unknown time. For four experiments, the sampling duration was not reported. The most used analysis method was LC-MS (54%), followed by GC-MS (31%), while other and mixed approaches were used less (8 and 7%, respectively).

**Figure 4:**
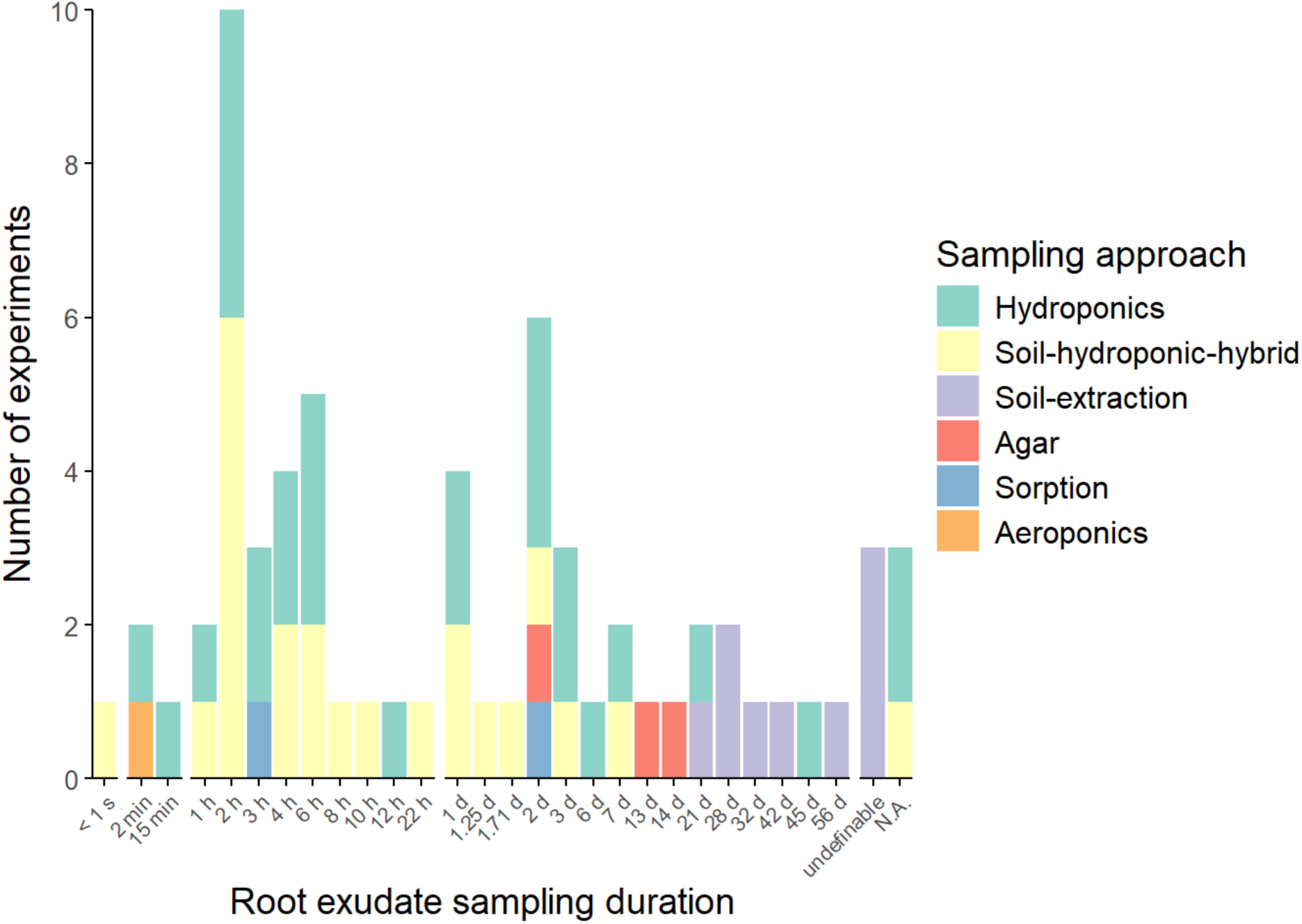
Contribution of different root exudate sampling durations and sampling approaches to the total number of experiments. Sampling durations ranged from less than one second to several weeks. Undefinable indicates that it is unknown how long root exudates have accumulated in the soil, while N.A. means that the studies did not report their sampling duration.

### Metabolic composition of root exudates

On average, each plant exuded 960 metabolites (7–33,870, median: 150). Of these metabolites, less than 5% (mostly in metabolic diverse exudates with >100 metabolites) or more than 95% (mostly in metabolic poor exudates with ≤100 metabolites) were annotated in each a quarter of all studies (Figure 5 A, B and C). By trend, GC-MS and mixed analyses resulted in fewer total metabolites being reported (9–1110, median: 56 (GC-MS); 31–464, median: 49 (mixed)), of which a higher percentage was annotated (24–100%, median: 73% (GC-MS) and 88% (mixed); Figure 5 B). In contrast, analyses with LC-MS resulted in more total metabolites (15–14,954, median: 200), of which only a fraction was annotated (29–100%, median: 17%; Figure 5 C).

**Figure 5:**
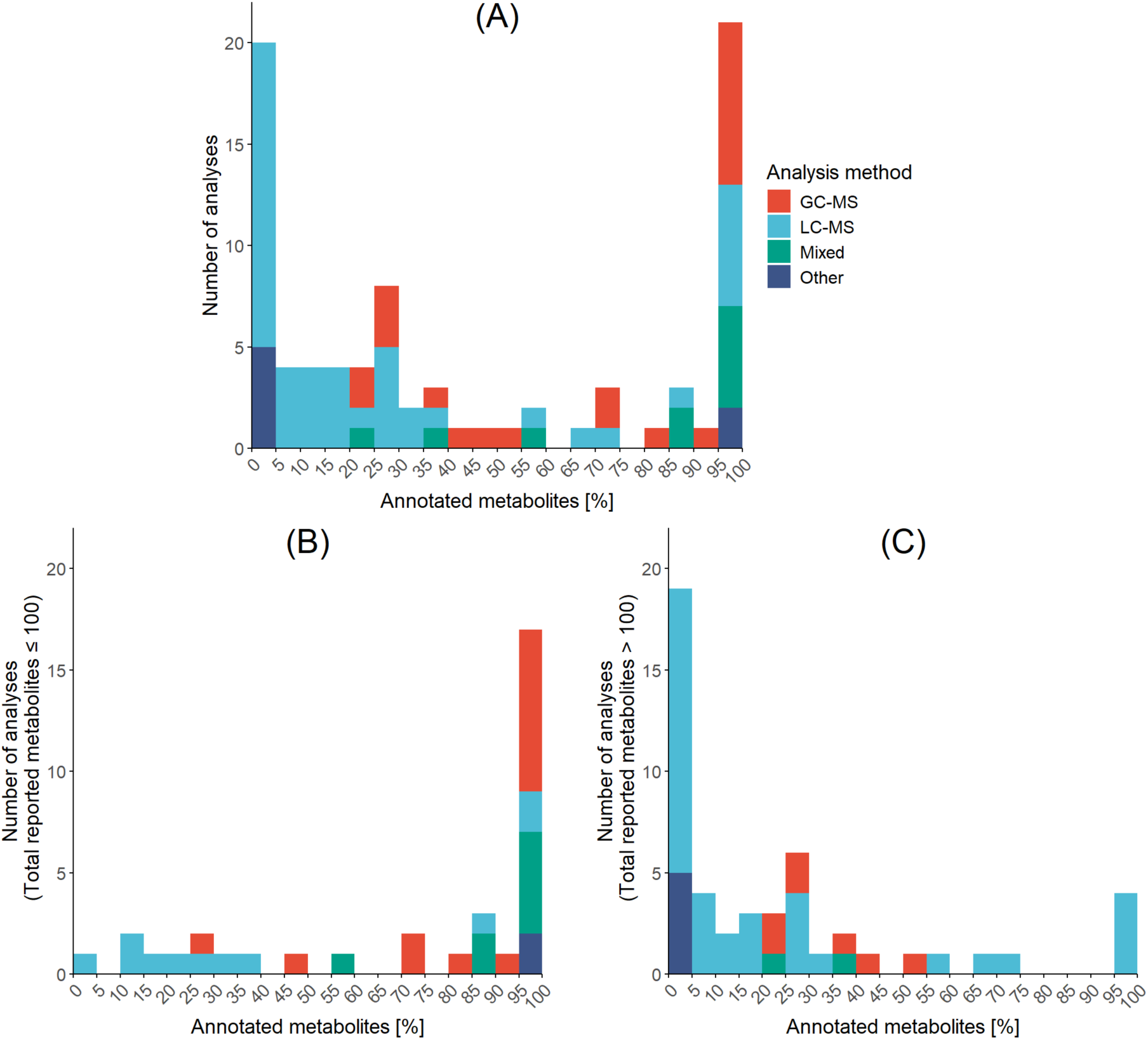
Frequency of percentage classes for annotated metabolites identified from root exudates by different analytical methods. (A) Includes all publications available, (B) includes percentage classes for annotated metabolites identified from metabolic poor (≤100 metabolites) exudates and (C) metabolic diverse (>100 metabolites) exudates. Mixed analysis methods use more than one analysis method, including GC-MS and/or LC-MS. Other methods include FT-ICR-MS and NMR.

Across all analyses, a slight majority of the annotated compounds of root exudates were shikimates and phenylpropanoids, and carbohydrates, followed by amino acids and peptides, and fatty acids. Polyketides, terpenoids and alkaloids had a smaller portion in the annotated root exudate metabolites (Figure 6, Supplementary Table 3). The identified compound classes differed among the analytical methods. When root exudates were analysed by GC-MS, carbohydrates (primary compounds) covered the largest portion (21%) of the annotated metabolites. When root exudates were analysed by LC-MS, shikimates and phenylpropanoids (secondary compounds) had the largest portion in the annotated metabolites (25%).

**Figure 6:**
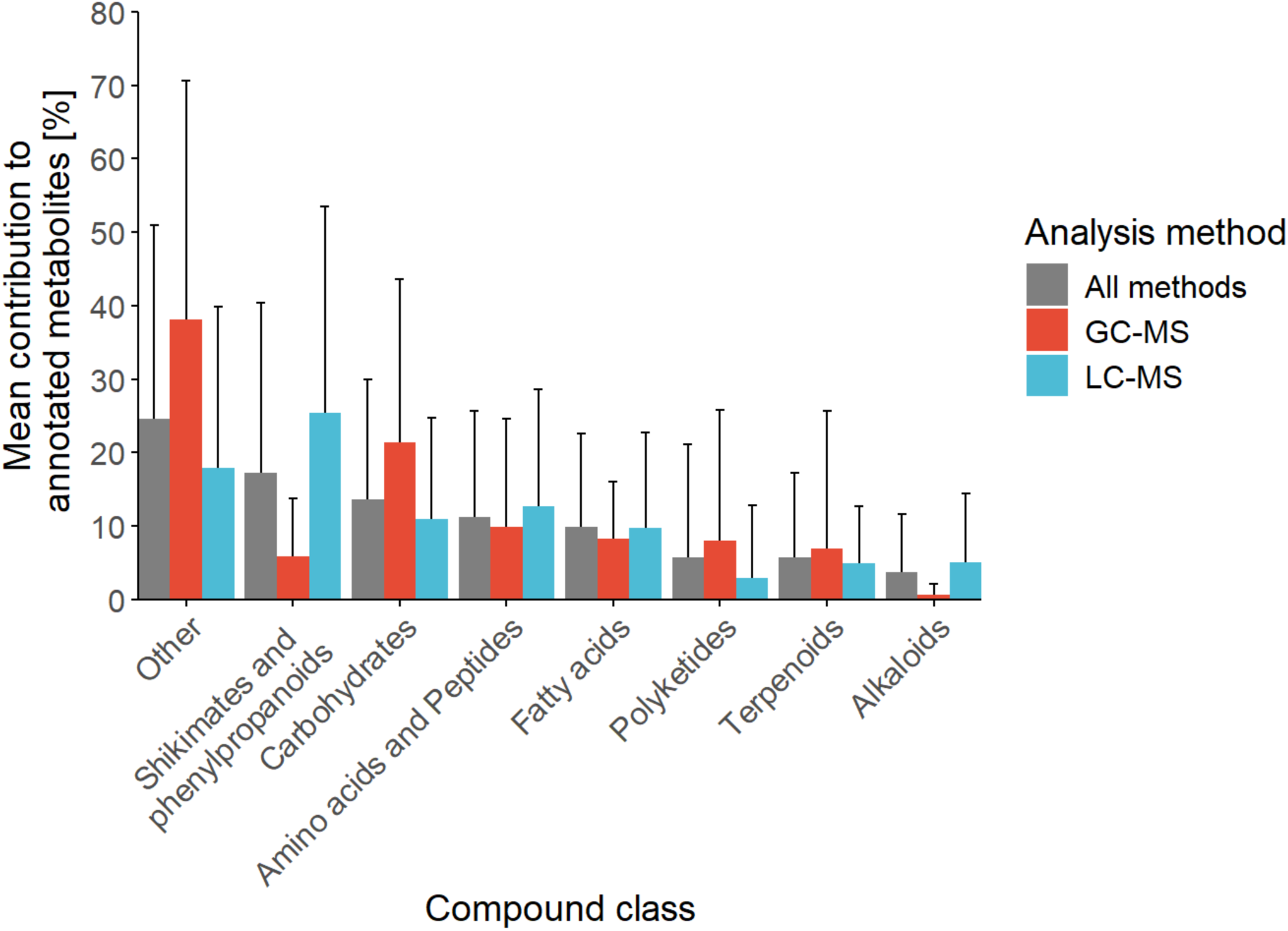
Composition of analysed root exudates with respect to the affiliation of annotated metabolites to substance classes. ‘All methods’ include GC-MS, LC-MS, mixed analyses and other analyses (mixed and other analyses are not shown separately because of their low abundance). The high standard deviation in the data results from many studies in which one or more compound classes are absent from the annotated data.

## Discussion

We identified several limitations in the studies, which will be addressed in this section. The limitations can be categorised into five sections: methodological limitations, comparability, underrepresentation of untreated samples, the gap in studies covering the whole lifetime of plants and the lack of chemical annotation coupled with the analysis of ecological function.

Firstly, studies are restricted by methodological limitations. There is a bias when deciding on an approach for root exudate sampling. Most studies we examined used the hydroponic approach (Figure 3); some combined it with a sterile system. On the one hand, only sterile systems can provide information on the original exudate composition because, in non-sterile conditions, microorganisms quickly process or degrade exudates (Kuijken et al., 2015). On the other hand, those sterile systems are far from resembling natural conditions and do not display effects caused by plant-microbe interactions. Also, the chemical composition of exudates in hydroponics and soil-grown plants differ, even if their compound classes are similar (Heuermann et al., 2023). Soil-extraction approaches can lead to the underrepresentation of some metabolites due to adsorption to soil particles. Furthermore, microbe-derived and microbial-processed metabolites are also included, which hampers the comparability with exudates sampled from hydroponics. The second most common approach was a soil-hydroponic hybrid, which combines the advantages of both approaches and might be the most promising sampling approach. However, chemical changes in the exudates can also happen after sampling due to the reaction of metabolites with each other (Lu et al., 2017). Further, caution also needs to be taken regarding mass spectrometry peak misidentification caused by isomers, overlapping compounds, and in-source degradation products (Alseekh et al., 2021). Also, peak resolution and detection thresholds of the chosen analytical platform for mass spectrometry influence which metabolites are found (Lu et al., 2017).

Secondly, comparing results across different studies is challenging. Variations in the sampling strategy and sample processing can lead to results that are not directly comparable (Freschet et al., 2021). Conditions influencing the exudate metabolome profiling results include, among others, the sampling solution used, the recovery time, sampling duration, time of day, plant age, and sterility of the sampling environment (Döll et al., 2024; Oburger & Jones, 2018). Because those conditions cannot be held constant between all studies targeting different research questions, and many protocols for exudate collection exist (Figures 3 and 4), they should at least be reported in detail. Also, we highly encourage researchers to include different conditions (such as different sampling durations or plant ages) to make studies more comparable. Our knowledge of exudate metabolomes is mainly based on crops, of which most are annual plants, and many belong to the Poales. Plants from natural ecosystems, such as trees and non-agricultural wetland plants, were underrepresented in studies on the exudate metabolome (Figure 2), leaving the variation of the exudate metabolome of non-agricultural plants understudied. This can bias the understanding of root exudates, mainly because crops grow under entirely different environmental conditions. Also, various plants differ in their adaption to the environment, life strategy and plant-internal C cycling. Different analysis methods (such as GC-MS or LC-MS) have advantages and shortcomings and influence the result (Lu et al., 2017). For higher comparability, details of the metabolite analysis should be described. This includes, for example, the metabolite class, molecular formula and identification level, as proposed by Sumner et al. (2007) and Alseekh et al. (2021). When naming the annotated metabolites, they should be identifiable by providing information on the structure. Also, the database used should be mentioned since different databases can lead to different results (Koistinen et al., 2023). A standard classification system, like the one advocated by Kim et al. (2021) and implemented, for example, by Döll et al. (2024) can also boost the comparability of untargeted studies of the exudate metabolome. The comparability should also be considered if the exudate composition is not the focus of a study but only a part of it, for example, in studies on the root microbiome or exudate addition experiments.

Thirdly, there is an underrepresentation of studies reporting on untreated samples since most studies focus on the effects of environmental factors like soil type, microbiome interactions, and plant stress conditions on root exudation profiles. More studies on natural variation in multiple genotypes, cultivars or populations under the same environmental influences are required to grasp the variability of root exudates in different species fully. If a study focuses on environmental effects, the metabolome of the control should also be reported instead of reporting only the differences between the control and treated samples.

Fourthly, there is a gap in studies spanning the total plant development. Most studies analyse exudation at a single time point in the plant’s life, mostly in young plants. This can lead to a bias of what is seen as the typical root exudate metabolome because exudate composition has been found to vary between different life stages of annual plants (Aulakh et al., 2001; Chaparro et al., 2013; Dechassa & Schenk, 2004; Santangeli et al., 2024) and between roots of different ages in trees (Michalet et al., 2013). In the best case, multiple growth stages or differently aged tissues should be studied. While this is possible for annual plants, it is challenging for perennial plants. In this case, plant or tissue age should at least be reported.

Lastly, there is a need to improve the chemical annotation of detected metabolites to link chemical composition to ecological functions such as plant-microbe interactions, allelochemical interactions, defence against pathogens, reaction to abiotic stressors, plant growth or organic matter decomposition. While many analysed compounds are still completely unclassified (Figure 5), most compounds’ ecological function is unknown. For this, the chemical composition of metabolites must be further verified under different conditions. Combining exudate metabolome analyses with genetic analysis of various cultivars and mutants can bring us one step further to finding the base exudate metabolome of specific plants (van Dam & Bouwmeester, 2016). Analysing the biological significance of root exudates is challenging because of the complex interactions of metabolites. Still, starting from this base knowledge, we can build up our understanding of exudate composition and functions under specific stressors, environmental influences, and biotic interactions. Exudate addition experiments can further enhance the knowledge of the ecological function of exudates, showing the direct effect of exudates or exudate compounds on the rhizosphere.

The abovementioned limitations might bias our understanding of the ecological functions of root exudate composition. Still, the increasing number of studies analysing the root exudate composition and environmental impact, pointing to a rise of technical opportunities and rising interest in untargeted metabolomics of root exudates, are promising (Figure 1). While it appears impossible to develop a sampling method for exudate collections that avoids all technical constraints, detailed reporting can significantly improve the comparability of studies. Also, closing the mentioned technical and knowledge gaps can boost our understanding of root exudates’ composition and function.

### Future directions

To stride ahead towards understanding the functional relevance of root exudates, we propose a framework that includes four steps (Figure 7). (1) The first crucial step is the sampling of representative exudates. For this, sampling from soil-grown plants (e.g. soil-hydroponic-hybrid) can be compared with sampling from sterile hydroponics. The combination would make it possible to sample ecologically relevant exudates from soil-grown plants while also sampling exudates unaltered by microorganisms for higher comparability between studies. Increased knowledge of differences between exudates from plants grown in soil and sterile hydroponics can also help put studies in context where only either one of the regarding the analytical platform. Direct comparison of GC-MS and LC-MS analysis of root exudates will allow us to get a broader picture of the composition of exudates. Combining both methods will also help to broaden the spectrum of captured components because different compound classes are better captured by different methods (Figure 6). Depending on the research question, untargeted approaches can be coupled with targeted approaches for key metabolites or other techniques such as NMR or FT-ICR-MS. (3) In the third step, advanced analytical techniques and bioinformatic tools need to be developed to enhance metabolite identification, quantification, and data integration. Those tools can include machine learning to improve the process. (4) Lastly, the exudate metabolomes need to be combined with multi-omic techniques and eco-evolutionary data to build a comprehensive understanding of the root exudation process. Multi-omics can include genomics, transcriptomics, and proteomics, as well as metabolomics from plant tissues and phloem sap. Combining the results with eco-evolutionary data will deepen our knowledge of the chemical diversity of root exudates and their role in adaptation. (+) Additionally, if possible, studies can be conducted on different genotypes, cultivars, or populations from various environmental conditions to determine the natural variation of exudates and include multiple life stages or plant ages.

**Figure 7:**
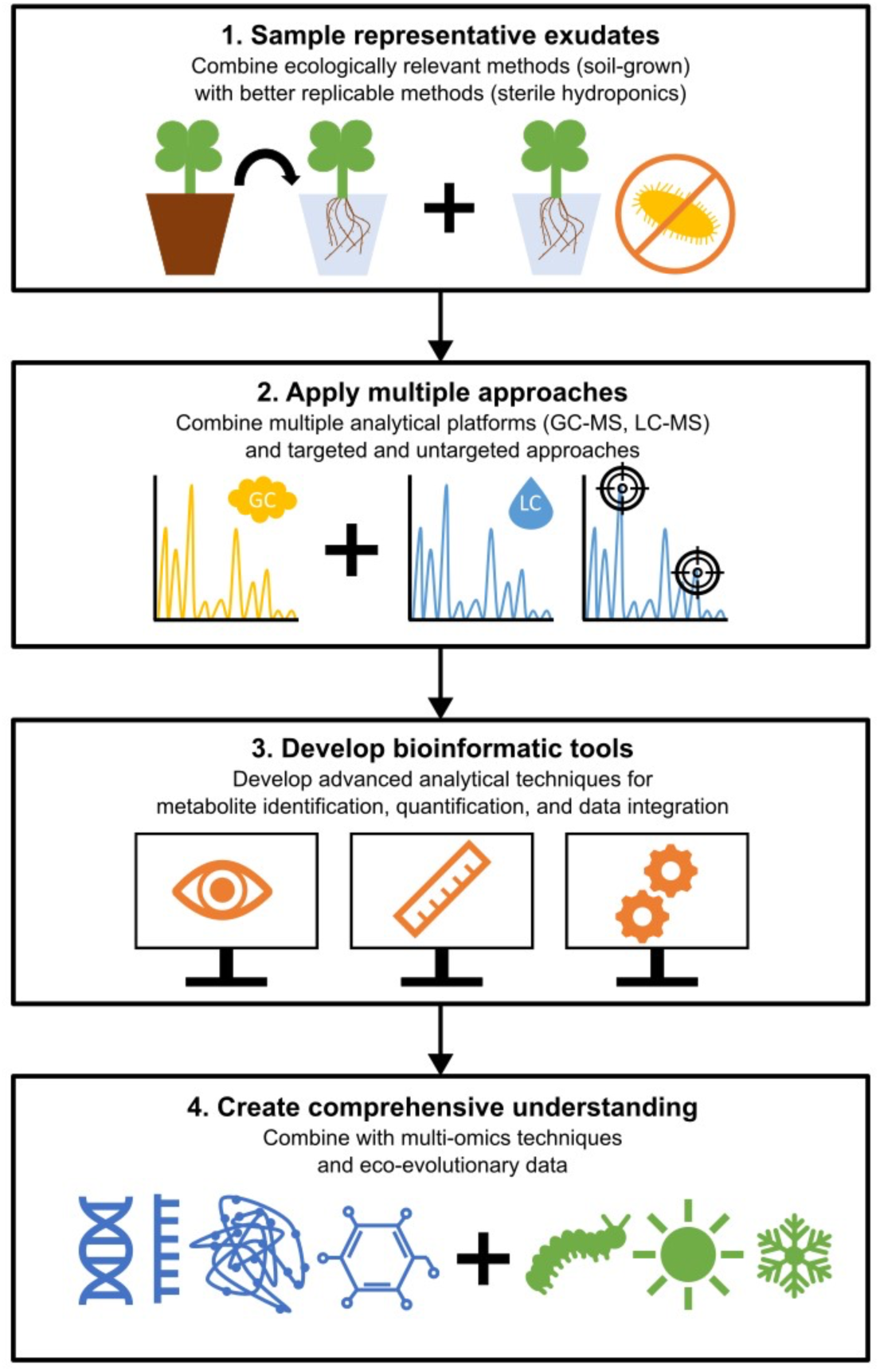
Framework for root exudate metabolomic investigations. A four-step framework to bring research closer to uncovering the chemical complexity of root exudates, including 1. Sampling of representative exudates, 2. applying multiple approaches, 3. Development of bioinformatic tools and 4. Creation of a comprehensive understanding.

Sampling root exudates is methodically challenging, and there are still gaps in the research on untargeted metabolomics of root exudates. However, the knowledge of root exudates is exponentially growing, bringing research closer to uncovering the chemical complexity of root exudates. Technological advances in LC-MS and GC-MS, making untargeted exudate analysis accessible to a broader community, and the recent developments in bioinformatics, machine learning, and AI enable researchers to annotate more root exudate metabolites. We advocate for obtaining root exudate data through well-defined sampling approaches followed by high throughput analytical and bioinformatic platforms. Integrating it with multi-omic techniques and eco-evolutionary data will provide a more comprehensive and multidisciplinary understanding of their chemical complexity and ecological functions.

## Supporting information

Metadata of papers used for analysis

## Authors contributions

HJS, KJ, and ICM conceived and designed the study. KM, AR, PJS and HJS contributed to developing the methodology and analysis framework. The manuscript was written by KM, AR, and PJS, with contributions from all authors. KM, AR, and PJS contributed to the data visualisation, including the creation of figures and graphs. All authors reviewed and approved the final manuscript.

## Conflict of Interest

The authors declare no conflict of interests.

---

**Table S1:** Excel table of our results of the literature search.

**Table S1:**
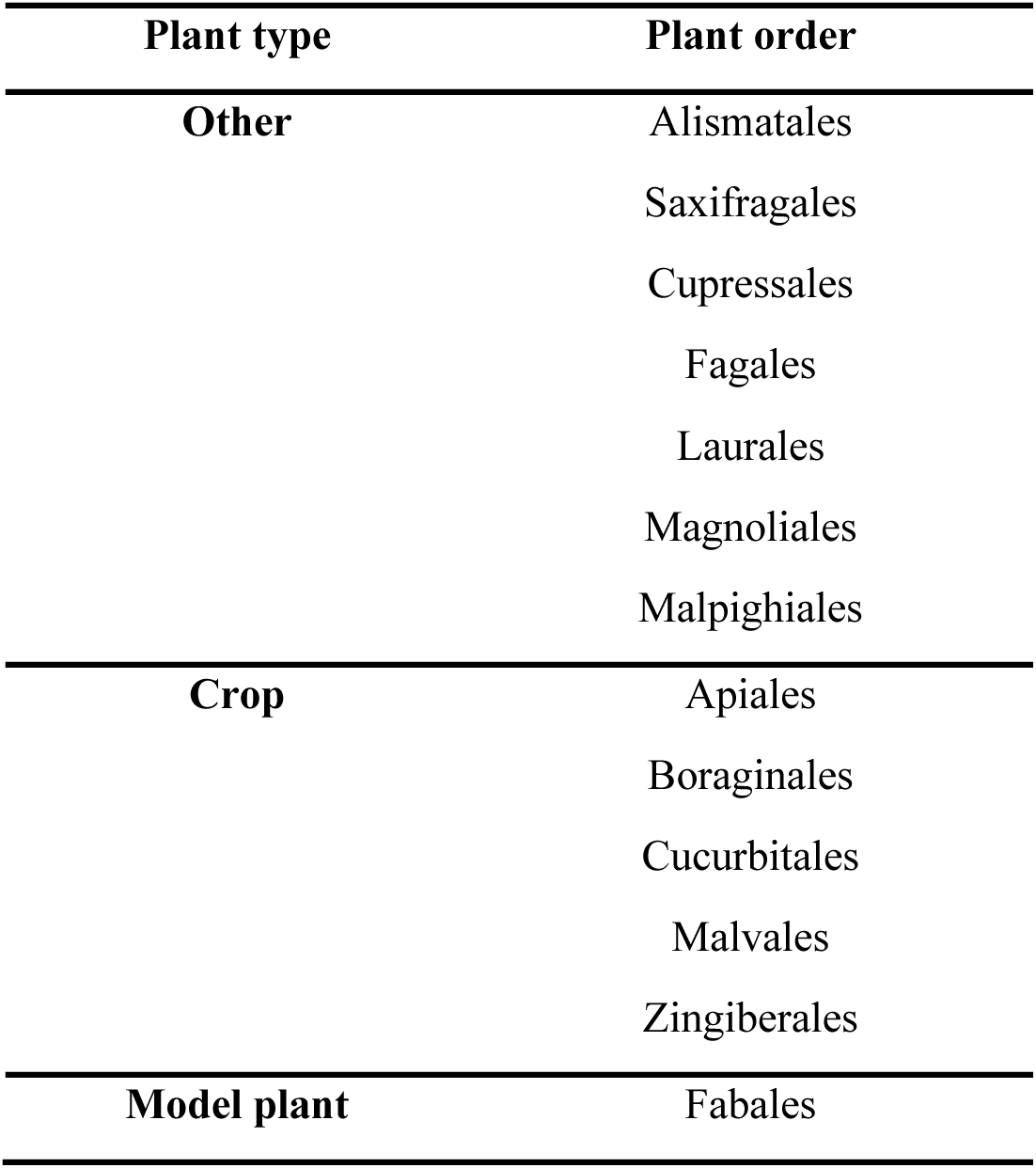
Plant orders grouped into the category ‘Other’ in Fig. 2.

**Table S2:** Mean, standard deviation and median of the contribution of each compound class to the total annotated compounds in per cent.

| Analysis method | Compound class | Mean [%] | Standard deviation [%] | Median [%] |
| --- | --- | --- | --- | --- |
| **GC-MS** | Alkaloids | 0.65 | 1.36 | 0 |
|  | Amino acids | 9.86 | 2.69 | 6.2 |
|  | Carbohydrates | 21.36 | 22.18 | 17.5 |
|  | Fatty acids | 8.2 | 7.95 | 3.43 |
|  | Polyketides | 8 | 17.73 | 0.43 |
|  | Shikimates and phenylpropanoids | 5.83 | 7.95 | 3.84 |
|  | Terpenoids | 6.86 | 18.76 | 0.45 |
|  | Others | 38.16 | 15.9 | 38.99 |
| **LC-MS** | Alkaloids | 5.04 | 9.39 | 0 |
|  | Amino acids | 12.68 | 15.94 | 0 |
|  | Carbohydrates | 10.91 | 13.77 | 7.33 |
|  | Fatty acids | 9.68 | 9.57 | 6.93 |
|  | Polyketides | 2.93 | 9.9 | 0 |
|  | Shikimates and phenylpropanoids | 25.37 | 28.14 | 12.55 |
|  | Terpenoids | 4.96 | 7.66 | 0 |
|  | Others | 17.95 | 15.9 | 10.49 |
| **All methods** | Alkaloids | 3.64 | 7.99 | 0 |
|  | Amino acids | 11.25 | 14.37 | 5 |
|  | Carbohydrates | 13.64 | 19.84 | 5.3 |
|  | Fatty acids | 9.9 | 10.97 | 7.98 |
|  | Polyketides | 5.71 | 11.23 | 0 |
|  | Shikimates and phenylpropanoids | 17.24 | 23.09 | 0 |
|  | Terpenoids | 5.64 | 11.59 | 0 |
|  | Others | 24.54 | 22.64 | 18.9 |

## Notes

### Competing Interest Statement

The authors have declared no competing interest.

